# VINYL: Variant prIoritizatioN bY survivaL analysis

**DOI:** 10.1101/2020.01.23.917229

**Authors:** Matteo Chiara, Pietro Mandreoli, Marco Antonio Tangaro, Anna Maria D’Erchia, Sandro Sorrentino, Cinzia Forleo, David S. Horner, Federico Zambelli, Graziano Pesole

## Abstract

**Motivation:** Clinical applications of genome re-sequencing technologies typically generate large amounts of data that need to be carefully annotated and interpreted to identify genetic variants associated with pathological conditions. In this context, accurate and reproducible methods for the functional annotation and prioritization of genetic variants are of fundamental importance, especially when large volumes of data - like those produced by modern sequencing technologies - are involved.

**Results:** In this paper, we present VINYL, a highly accurate and fully automated system for the functional annotation and prioritization of genetic variants in large scale clinical studies. Extensive analyses of both real and simulated datasets suggest that VINYL show higher accuracy and sensitivity when compared to equivalent state of the art methods, allowing the rapid and systematic identification of potentially pathogenic variants in different experimental settings.

## Introduction

Applications of modern high throughput genome sequencing technologies to healthcare and clinical practice are driving a major breakthrough in medical science (Saudi Mendeliome Group 2015, UK10K Consortium 2015, Kowalski et al 2019). The unprecedented ability to interrogate the (more than) 3 billion pairs of nucleotides that compose our genome, in a systematic and reliable manner, provides a formidable tool for the characterization and functional annotation of the human variome, that is, the complete set of genetic variants found in the human population (Gurdasani et al 2015, Kowalski et al 2019, Nagasaki et al 2015). In these settings, the capacity to link genetic variants with phenotypic traits, pathological conditions, and/or positive or adverse reactions to therapies and medications will be of instrumental importance for the development of informed approaches to medical science, such as precision medicine (Lu et al, 2014), that is the possibility for patients to be treated based on their genetic background, or predictive medicine (Kotze et al, 2015) where risk factors for various diseases can be accounted beforehand and suitable measures instituted in order to prevent the disease or mitigate its severity. Accordingly, numerous countries and institutions worldwide are already undertaking or are planning to launch large-scale projects aiming to sequence an increasing proportion of their population. These include, among the others, the UK10K project in the United Kingdom (UK10K Consortium et al, 2015), the All of Us research program by the NIH (All of Us Research program investigators, 2019), the French Plan for Genomic medicine funded by the French Ministry of Health (Lethimonnier and Levy, 2018), and the European ‘1+ Million Genomes’ initiative promoted by the European Community (Saunders et al 2019).

While the possibility to sequence an unprecedented number of human genomes could form the basis for a new revolution in medical science and human genetics, the need to handle, analyze and interpret large collections of “big” genomic data is posing major challenges to genomics and bioinformatics which at present remain unresolved (Alyass et al, Klein et al 2017, 2015, Horowitz et al, 2019, Stark et al 2019). These limitations are both technical, due to the need to develop dedicated infrastructures for the handling, sharing and processing of the data, and methodological, due to the need to integrate multiple bioinformatics tools and data formats into complex analysis workflows- which in turn require a substantial effort for their set up and optimization (Canzoneri et al 2019, Ginsburg and Phillips 2018, Servant et al 2014). More important, a typical Next Generation Sequencing assay can detect in the order of ten of thousands or even millions of genetic variants, all of which need to be carefully annotated and interpreted in order to identify genetic traits possibly associated with a pathological condition of interest (Elbeck et al 2017). This process, which is known as “variant prioritization” or “variant filtration”, typically requires manual curation by an expert clinician, and represents a major bottleneck for the application of large scale genotyping assays in clinical settings (Frebourg et al 2014, Jalali et al 2017). Moreover -although very detailed and rigorous guidelines for the interpretation and analysis of genetic variants in clinical settings are currently available- it is not uncommon for different institutions/operators to apply slightly different criteria and filters, thus limiting the overall reproducibility of the results of this type of analysis (Pabinger et al 2014).

In this paper, we present VINYL, a novel fully automated workflow for the prioritization of genetic variants in clinical studies. By building on guidelines and recommendations derived from clinical practice (Richards et al, 2015), VINYL derives a pathogenicity score by aggregating different sources of evidence and annotations obtained from publicly available resources for the functional annotation of human genetic variants. Several studies (Cirulli et al 2015, Lee et al 2014, Moutsianas et al 2015, Guo et al 2016, Li et al 2008) have reported that populations of affected individuals are expected to harbor an excess of deleterious or slightly deleterious variants at disease-associated loci with respect to a matched population of unaffected controls. VINYL uses this principle to optimize the computation of its composite pathogenicity score. An automatic procedure based on survival analysis is applied, wherein the optimal balance between the components of the scoring system as well as the ideal threshold for the identification of potentially pathogenic variants are established by identifying the scoring system that maximize the difference between the number of potentially pathogenic variants identified in a population of affected individuals and the equivalent figure from a population of matched controls. We perform extensive simulations based on publicly available human genetic data to demonstrate the validity of our approach, and test the ability of VINYL to detect different types of genetic variants associated with pathological conditions. Finally, we apply our tool to a cohort of 38 patients with a diagnosis of dilated cardiomyopathy (DCM), arrhythmogenic right ventricular cardiomyopathy (ARVC) or hypertrophic cardiomyopathy (HCM) who were previously subjected to genotyping by targeted resequencing of a panel of 115 genes associated with cardiomyopathies/channelopathies (Forleo et al, 2017). We show that, while attaining very low levels of false positives, VINYL compares favorably to other state of the art methods for the prioritization of genetic variants both on real and simulated data. More importantly, we demonstrate that our tool is capable to correctly identify all the variants that were previously classified as Pathogenic/Likely Pathogenic by careful expert manual curation on the Forleo et al dataset. All in all, we believe that by providing a rapid and systematic approach for the prioritization of genetic variants, VINYL can greatly facilitate the identification of pathogenic or potentially pathogenic variants in large scale clinical studies. VINYL is available at: beaconlab.it/VINYL. To facilitate the usage of the tool and to improve the reproducibility of the analyses, VINYL is incorporated into a dedicated instance of the popular Galaxy workflow manager (Afgan et al, 2018), along with a highly curated collection of tools and resources for the functional annotation of genetic variants.

## Methods

### Implementation of VINYL

VINYL is implemented as a Laniakea (Tangaro et al, 2018) Galaxy (Afgan et al, 2018) instance based on Galaxy release 18.05. Annotation of VCF files is performed by the Annovar software (Wang et al, 2010), using a collection of “standard” resources maintained by the Annovar developers along with a selection of custom annotation tracks. These include the OregAnno database (Griffith et al 2008), the Ensembl regulatory build annotation (Zerbino et al 2016), the NHGRI-EBI GWAS catalog (Buniello et al, 2019) and the ncER score, which provide fine-grained annotations of non-coding and regulatory genomic elements (Wells et al 2019). A complete list of the annotation tracks that are currently supported by VINYL along with a brief description is reported in Supplementary Table S1. The VINYL application itself is implemented as a collection of Perl and R scripts and is composed of 3 main modules:

- *the optimizer*, which computes the optimal weights for the components of the pathogenicity score by performing a grid search over the parameter space;
- *the threshold optimizer*, that derives the optimal score threshold for the identification of likely pathogenic variants
- and the *score calculator*, the main tool which computes the pathogenicity scores.

These tools can be executed independently, or via an automated workflow which is available in the VINYL Galaxy instance. All the software is currently available from http://beaconlab.it/VINYL. A detailed manual for the usage of VINYL is available at http://90.147.75.93/galaxy/static/manual/. VINYL is available as a standalone command line-tool from https://github.com/matteo14c/VINYL.

### Computation of the pathogenicity score

VINYL computes its pathogenicity score directly from an Annovar annotated VCF file. Annotations that should be considered for the computation of the score can be specified by a plain text configuration file. Currently VINYL can discriminate between 11 different types of functional annotations, including -among the others- databases of human genetic variation (RV), the predicted functional effects of the variants (FE) and/or their presence/absence in databases of clinically relevant genetic variants (DB). A complete list is reported in Supplementary Table S2 (and in the online manual). The score itself is computed as an aggregated score that combines all the different types of functional annotation by the means of a simple linear formula:

## Pat Score= DB+RV+FE+NS+OR+eQ+AD+mi+Reg+TF+GW+Sp

### Single components of the score are computed according to the following rules

- **DBs of pathogenic variants (DB)**: the score is incremented if variants are described as Pathogenic or Likely Pathogenic in publicly available resources of clinically relevant variants. The score is decreased for variants that are reported as “Benign” or “Likely Benign”. Users can provide a description of the disease and its symptoms using a simple configuration file. Only entries that match these keywords are considered for the computation on the score. In the current implementation of VINYL the Clinvar (Landrum et al, 2014) database is used as the main source for the annotation of disease-associated genetic variants
- **Rare Variants (RV):** the score is increased if a genetic variant shows a Minor Allele Frequency (MAF) lower than a user-defined cutoff -typically the prevalence of the disease- in public databases of human genetic variation
- **Functional effect of the variant (FE)**: the score is increased if the variant is predicted to have a deleterious functional effect (splicing variants, stop-gain, frameshift variants).
- **Disruptive non-synonymous (NS):** the score is incremented for non-synonymous variants that are predicted to have a disruptive effect. Tools to be considered for the evaluation of the effect of NS variants can be specified at runtime. Predictions are derived from the dbNFSP database (Liu et al, 2016) version 3.5a.
- **Overrepresentation (OR):** if a genetic variant with MAF ≤0.01 (the frequency cut-off that is normally considered for the definition of “common” Single Nucleotide Polymorphism) is found in N or more affected individuals the score is incremented. The value of N is specified at runtime, or set to 10% of the size of the cohort otherwise.
- **eQTLs (eQ):** when a variant is associated with an eQTL according to the GTEx study (GTEx Consortium, 2013) the score is incremented. A list of relevant (to the pathological condition) tissues for the annotation of eQTLs (according to the GTEx nomenclature) can be provided by users in the form of a simple text file.
- **Disease-associated genes (AD):** the score is incremented if a genetic variant is associated with genes previously implicated in the disease or in similar pathological conditions. Users can provide a list of disease-related genes by means of a simple configuration file
- **miRNA binding site (mi):** the score is increased if the variant is associated with a known miRNA binding site
- **Regulatory element (Reg):** the score is incremented if the variant is part of a genomic regulatory element (promoter, enhancer, silencer), according to the OregAnno database (Griffith et al 2008), the Ensembl regulatory build annotation (Zerbino et al 2016)
- **TF binding site (TF):** the score is increased if the variant is associated with a transcription factor binding site, according to the OregAnno database (Griffith et al 2008), the Ensembl regulatory build annotation (Zerbino et al 2016)
- **GWAS (GW):** the score is incremented if the variant is associated with a phenotypic trait relevant for the pathological condition according to one or more GWAS studies. Similar to the DB score, only entries matching a user-specified list of keywords are considered for the computation of this score
- **Splicing variants (Sp):** the score is incremented if the variant is reported to have a deleterious effect on a splice site according to the dbscsnv11 (Jian et al, 2014) database

Additionally, users can configure the behavior of VINYL by providing configuration files and parameters, to specify a disease model (Autosomic Dominant, Autosomic Recessive or X-linked), a list of symptoms associated with the disease for a more accurate evaluation of the entries reported in publicly available databases (DB and GW scores), or to define a set of genes implicated with the pathological condition of interest and/or tissues to be considered for the evaluation of expression quantitative trait loci (eQTLs), to be used for the computation of the AD ad eQ scores respectively. When a disease model is specified, only genetic variants that are inherited according to that model are considered for the computation of the score.

### Optimization of the pathogenicity score

Genetic algorithms, as implemented in the genalg (Willighagen and Ballings, 2015) R library, are used to identify optimal weights for the components of the pathogenicity score by performing a search on the parameter space. Score distributions are computed for a population of affected individuals (A) and a population of healthy controls (C). The optimal scoring system and the corresponding threshold for the identification of potentially pathogenic variants are established by an iterative survival analysis based on the Wang Allison method (Wang et al, 2004). For every scoring system, possible cut-off values spanning from the maximum score to the minimum score with an interval of 0.5 are evaluated, and the number of predicted potentially pathogenic variants identified in the A and C populations are recorded. A Fisher’s Exact test is used to test the significance of the over-representation of likely pathogenic variants in A with respect to C. Finally, the scoring system (and the corresponding threshold value) that maximizes the difference between the number of potentially pathogenic variants identified in the population of affected individuals with respect to the control population, and that, at the same time, minimizes the number of potentially pathogenic variants identified in the control population is selected. The following equation is used to define the optimality criterion:

### Optimal Score=argmax< 0.5*-log10 (Fpv) +0.3*(Ffc) -0.2 *PC >

Fpv= p-value for the over-representation of likely pathogenic variants in A according to the Fisher’s Exact Test. Ffc= ratio between the proportion of likely pathogenic variants identified in A and C respectively. PC= number of potentially pathogenic variants identified in C. Coefficients of the equation have been derived empirically, to obtain a reasonable balance between the maximization of the number of potentially pathogenic variants identified in A, and the minimization of the, likely false positive, pathogenic variants identified in C. Values can be modified by the users at runtime. Of notice, in all the experimental settings tested in this work no detectable changes in the sensitivity and specificity of VINYL were observed when different values were applied, suggesting that the genetic algorithms used for the optimization of the VINYL pathogenicity score can robustly converge to the optimal solution, irrespective of slight variations in the formulation of the optimization function.

### Utilities for the post-processing of VINYL’s output

All the utilities for the post-processing of VINYL’s output files are implemented in the form of standalone R scripts. Principal Component Analysis is performed by the means of the prcomp R function from the stats package (R Core Team 2018). Graphical representation of the results is obtained by the means of the R ggplot2 package (Wickham 2016).

### Variant calling and simulation of disease-causing variants

In the present study, variant calling was performed by the CoVaCS pipeline (Chiara et al 2018). Simulation of disease-causing genetic variants was performed by the means of the Hapgen2 (Su et al 2011) program using the haplotype files of the TSI (Toscani in Italia) population from the 1000G study to provide the genetic background.

The latest version of Hapgen2 was obtained from: https://mathgen.stats.ox.ac.uk/genetics_software/hapgen/hapgen2.html, while haplotype files from the 1000G project were obtained from: https://mathgen.stats.ox.ac.uk/impute/impute_v1.html#Using_IMPUTE_with_the_HapMap_Data. To simulate different levels of association with pathological conditions, three different distributions of Odds Risk Ratios, with an average Risk Ratio of 3,10 and 20 respectively, were simulated using the rnorm function in R. Standard deviation was set to 10% of the average.

Cohorts of different size, formed by 25, 50 and 100 individuals respectively were simulated, including a variable number of polymorphic positions: 1000, 5000 and 10000, to simulate different sequencing strategies. Disease-associated variants have been simulated by randomly selecting a matched number of rare variants (Minor Allele Frequency ≤0.001) with different predicted functional effects, including splice site variants, variants in promoter regions, frameshift variants, variants in miRNA target regions, stop-gain/stop-loss variants. At every iteration, a total number of 75 distinct disease causing variants were simulated, of which a maximum of 10 were implicated in a known pathological condition according to ClinVar.

In the analysis of the Forleo et al (2017) dataset the TSI (Toscani in Italia) population from the 1000G (The 1000 Genomes Project Consortium, 2015) study was used as the “control” population.

### Execution of Privar and KGGseq

The latest versions of KGGseq (Li et al 2012) and Privar (Zhang et al 2013) were obtained from http://grass.cgs.hku.hk/limx/kggseq/ and http://paed.hku.hk/genome/software.html, respectively. Privar was executed using the “Literature-based strategy” with default parameters. A custom list of disease-associated genes (identical to that used for VINYL) was provided by the means of the “- customlist” parameter.

KGGseq was applied using the strategy illustrated in the reference manual for the prioritization of genetic variants associated with rare Mendelian diseases; “the “--candi-list” and “--phenotype-term” parameters were used to provide a list of disease-associated genes and a list of symptoms of the disease under study, respectively. Both lists were completely identical to the lists used in VINYL to provide the same type of information. Consistent with the parameters used in VINYL, the cut-off frequency for rare alleles was set to 10e-4.

## Results

### VINYL: an automated tool for variant prioritization

VINYL provides a fully automated system for the prioritization of pathogenic variants, which -similar to the guidelines used in clinical practice- includes a scoring system based on the integration of different types of annotations. By leveraging the Galaxy workflow manager, VINYL is made available through a powerful and user-friendly web-based graphical interface and allows collaborative and highly reproducible analysis of large amounts of data. Encrypted data volumes are used to ensure data protection. Users can upload their data to VINYL in the form of plain VCF files. Variants annotation is performed by the Annovar software (Wang et al, 2010), which is available in VINYL along with an extensive collection of resources for the annotation of genetic variants (see Table S1). Additional information, which is used for the computation of the pathogenicity score, including for example the symptoms and prevalence of the pathological condition under study, the model of inheritance of the disease, the types of predicted functional effects that should be considered deleterious, the specific computational tools to be used for the prediction of the functional effects of genetic variants, can be specified by users at run-time using simple configuration files in plain text format (see Material and Methods, and Supplementary Materials).

The main output consists of a tabular file, where variants are ranked according to their pathogenicity score and the threshold for the identification of pathogenic variants is derived automatically. Additional utilities (see below) can be used to perform more fine-grained analyses for the identification of genes that display a significant over-representation of high scoring variants, or for the stratification of patients in groups by dimensionality reduction techniques. Moreover, along with a carefully curated collection of tools and resources for the functional annotation of genetic variants, the VINYL Galaxy instance incorporates also a collection of reference data, including VCF files of 26 distinct human geographic populations from the 1000 Genomes study (The 1000 Genomes Project Consortium, 2015), which can provide a suitable background control population for most clinical studies. The features contained in VINYL and the rationale used in the implementation of the tool are briefly outlined in Figure 1.

**Figure 1:**
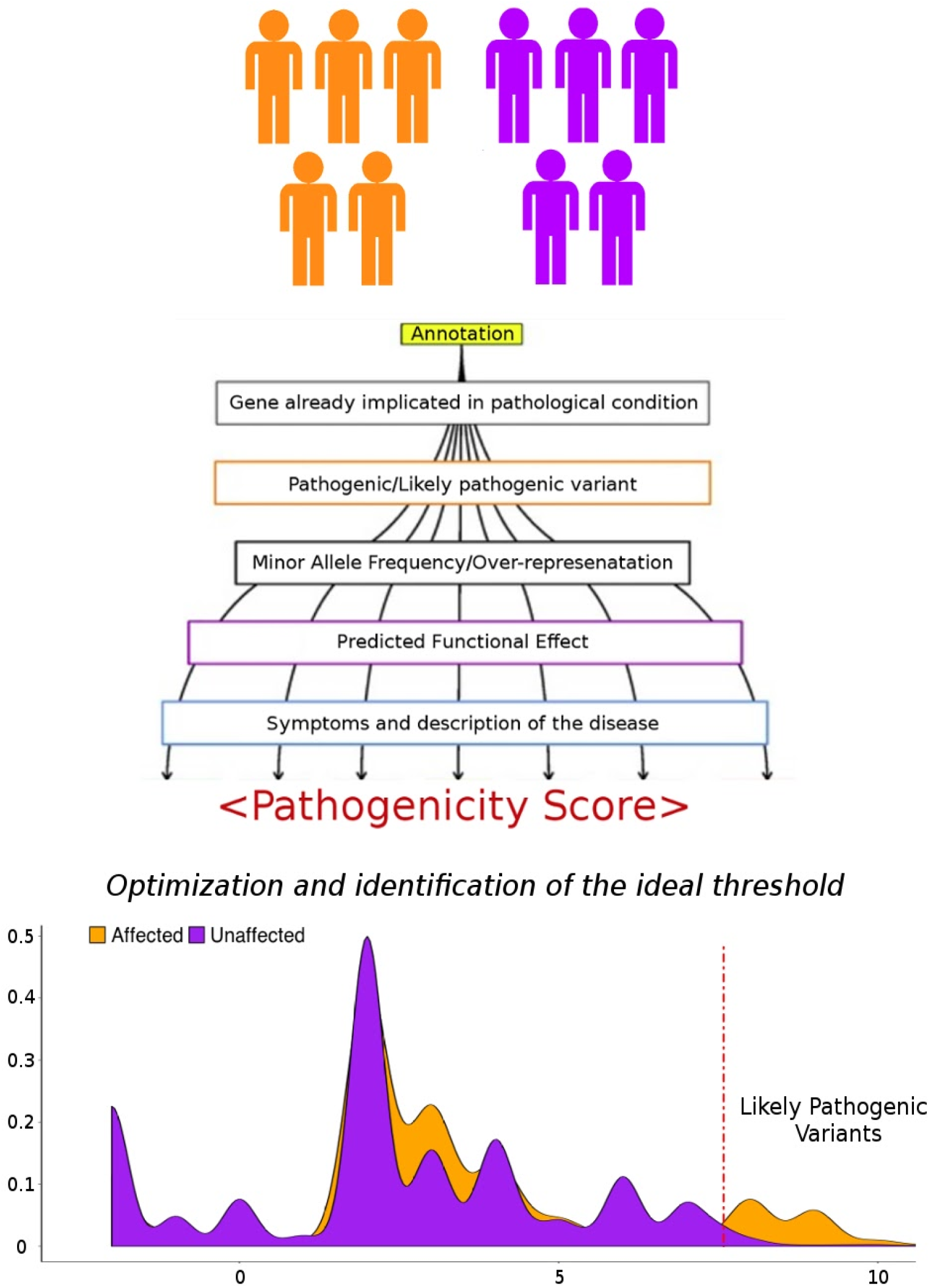
Outline of the variant prioritization strategy adopted by VINYL. Genetic variants identified from a cohort of affected individuals (orange) and a cohort of healthy controls (purple) are subjected to variant annotation. A scoring algorithm is subsequently used to compute a pathogenicity score based on the predicted functional effect of the variants. Different scoring schemes are evaluated and distributions of pathogenicity scores are compared between the 2 cohorts (affected and controls). The scoring system that maximizes the difference of the score distribution between the 2 population is selected. The corresponding cut-off score for the identification of potentially pathogenic variants is identified as the threshold that maximizes the number of potentially pathogenic variants in the cohort of affected individuals, while at the same time minimizing the number of potentially pathogenic variants in the control population.

### Evaluation of VINYL on simulated data

The ability of VINYL to correctly identify genetic variants associated with pathological conditions has been evaluated by performing extensive simulations of disease-associated variants derived from real human haplotypes and comparing the performances attained by our tool with two other popular methods for the prioritization of genetic variants: *Privar* (Zhang et al, 2013) and *KGGseq* (Li et al, 2012). Different scenarios were simulated in order to evaluate the impact of the analysis of cohorts of different size (25, 50 and 100 individuals), the number of disease-associated variants (20, 50 and 100), the strength of the association of these variants with the pathological condition (odd risk ratio of 5, 10 and 20) and the total number of variants included in the call-set (1000, 5000 and 10000), a proxy for evaluation of the usage of different sequencing strategies (from targeted resequencing of a limited number of genes to WES). Finally, to investigate the ability of the tools considered to correctly prioritize disease-associated variants of different types, a matched number of variants with different predicted functional effects were simulated (see Materials and Methods).

As outlined in Table 1 and Figure 2, we observe that, while achieving a remarkable level of accuracy, with a false positive rate that is consistently below 1%, VINYL demonstrates an improved sensitivity in the detection of disease-associated genetic variants compared to both KGGseq and Privar, resulting in a significantly increase in AUC (area under the curve) in all the simulations performed in this study. These results suggest that the approach adopted by VINYL can outperform currently available state of the art methods in the prioritization of disease-associated genetic variants. This notwithstanding, we notice that- as expected- the performances of VINYL are strongly influenced by the composition and the size of the input dataset, as we observe an increase in sensitivity when large cohorts of patients are analyzed and/or when the number of pathogenic variants included in the simulated cohort is increased (Table 1). Interestingly however, we notice that the size and breadth of the input dataset- both in terms of the number of individuals considered and number of genetic variants that are analysed- do not seem to have a major impact on the specificity of our tool, as we do not observe any detectable change in the False Discovery Rate when a higher number of variants and/or larger cohorts of individuals are considered. Since, unlike other similar methods, VINYL requires a population of negative controls to derive and optimize its variant prioritization system, we reasoned that the choice of the correct background population could have a major impact on the accuracy or our tool, thus limiting the applicability of VINYL to only cases when a control population with a similar genetic background is available. To test this hypothesis we repeated our analyses of simulated data by using 3 distantly related human geographic populations: CEU (Northern Europeans from Utah), CHB (Han Chinese) and ESN (Esan in Nigeria) from the 1000G study as a control. While we observe no detectable change in the accuracy of VINYL (Supplementary Table S3), we notice that the sensitivity of our method seems to be significantly decreased when a “mismatched” control population is used, and especially when distantly related populations are considered. Unsurprisingly, we notice that the importance of the scores related to allele frequency and over-representation of alleles in the population were systematically reduced in these settings, suggesting that reduction in sensitivity could be related to the presence of mildly deleterious polymorphisms showing a population biased frequency distribution. To test this hypothesis we repeated these analyses by excluding from the control population all the alleles with a minor allele frequency ≥0.01 and showing a difference in allele frequency of 4 fold or greater between any of the 26 geographic populations included in the 1000G study. When these population biased alleles are removed from the control population, a marked increase in sensitivity (Supplementary Table S4) is observed, suggesting that the effectiveness of the approach adopted by VINYL could be sensibly reduced in the presence of diffuse geographic allele frequency distribution biases. Accordingly, VCF files filtered from population biased alleles have been incorporated in the main Galaxy VINYL instance to serve as an alternative reference for studies where a geographically matched control population is not available.

**Table 1:** Sensitivity and specificity on simulated data. Levels of sensitivity and specificity of VINYL, Privar and KGGseq on simulated data. A) Dataset with 1000 polymorphic sites. B) Dataset with 5000 polymorphic sites. C) Dataset with 10000 polymorphic sites. Sizes of the simulated cohorts (25,50 or 100 individuals) are reported in the first column. Tools are indicated in the second column. Corresponding levels of sensitivity and specificity attained by each tool, are reported in the subsequent columns. Columns 3 to 4, 5 to 6 and 7 to 8, report the values for the simulation of pathogenic variants with an odd Risk Ratio of 3, 10 and 20 respectively.

| 1000 |  |  |  |  |  |  |  |
| --- | --- | --- | --- | --- | --- | --- | --- |
| A |  | RiskRatio~3 |  | RiskRatio~10 |  | RiskRatio~20 |  |
|  |  | <u>Sens</u> | <u>Spec</u> | <u>Sens</u> | <u>Spec</u> | <u>Sens</u> | <u>Spec</u> |
| 25 | VINYL | 73,01 | 99,66 | 78,89 | 99,29 | 89,95 | 99,69 |
|  | Privar | 54,97 | 93,44 | 59,79 | 91,64 | 61,58 | 93,57 |
|  | KGGSeq | 62,29 | 95,55 | 64,03 | 96,77 | 59,93 | 96,96 |
| 50 | VINYL | 78,64 | 99,52 | 86,14 | 100,36 | 94,35 | 99,72 |
|  | Privar | 56,66 | 94,01 | 58,60 | 90,04 | 59,95 | 90,79 |
|  | KGGSeq | 63,92 | 95,05 | 66,37 | 96,60 | 65,90 | 94,56 |
| 100 | VINYL | 82,69 | 99,89 | 90,47 | 99,04 | 96,27 | 99,82 |
|  | Privar | 55,80 | 89,87 | 58,06 | 91,34 | 59,63 | 91,77 |
|  | KGGSeq | 66,34 | 96,14 | 64,47 | 96,59 | 66,91 | 96,58 |
| 5000 |  |  |  |  |  |  |  |
| B |  | RiskRatio~3 |  | RiskRatio~10 |  | RiskRatio~20 |  |
|  |  | <u>Sens</u> | <u>Spec</u> | <u>Sens</u> | <u>Spec</u> | <u>Sens</u> | <u>Spec</u> |
| 25 | VINYL | 76,95 | 99,32 | 82,90 | 99,33 | 91,50 | 99,96 |
|  | Privar | 58,27 | 92,47 | 61,67 | 91,70 | 63,30 | 92,99 |
|  | KGGSeq | 63,21 | 95,55 | 64,52 | 95,93 | 60,60 | 96,13 |
| 50 | VINYL | 80,75 | 99,41 | 87,12 | 99,54 | 96,03 | 99,26 |
|  | Privar | 56,84 | 93,54 | 58,84 | 89,80 | 62,40 | 90,36 |
|  | KGGSeq | 63,68 | 94,42 | 69,78 | 95,94 | 67,35 | 94,10 |
| 100 | VINYL | 84,53 | 99,25 | 93,83 | 99,17 | 99,86 | 99,63 |
|  | Privar | 58,47 | 89,89 | 62,42 | 90,59 | 60,78 | 91,06 |
|  | KGGSeq | 66,86 | 95,47 | 66,53 | 96,27 | 67,71 | 96,36 |
| 10000 |  |  |  |  |  |  |  |
| C |  | RiskRatio~3 |  | RiskRatio~10 |  | RiskRatio~20 |  |
|  |  | <u>Sens</u> | <u>Spec</u> | <u>Sens</u> | <u>Spec</u> | <u>Sens</u> | <u>Spec</u> |
| 25 | VINYL | 78,41 | 99,48 | 84,01 | 99,40 | 92,23 | 99,66 |
|  | Privar | 62,70 | 92,60 | 64,17 | 91,87 | 64,53 | 93,11 |
|  | KGGSeq | 69,85 | 95,75 | 67,57 | 96,08 | 61,25 | 96,30 |
| 50 | VINYL | 82,95 | 99,55 | 88,44 | 99,63 | 98,29 | 99,45 |
|  | Privar | 59,84 | 93,69 | 62,06 | 89,87 | 65,02 | 90,52 |
|  | KGGSeq | 65,59 | 94,59 | 73,40 | 96,07 | 68,64 | 94,27 |
| 100 | VINYL | 88,38 | 99,42 | 97,32 | 99,39 | 99,29 | 99,85 |
|  | Privar | 61,53 | 89,91 | 65,48 | 90,77 | 64,08 | 91,12 |
|  | KGGSeq | 68,82 | 95,49 | 70,21 | 96,28 | 72,02 | 96,51 |

**Figure 2:**
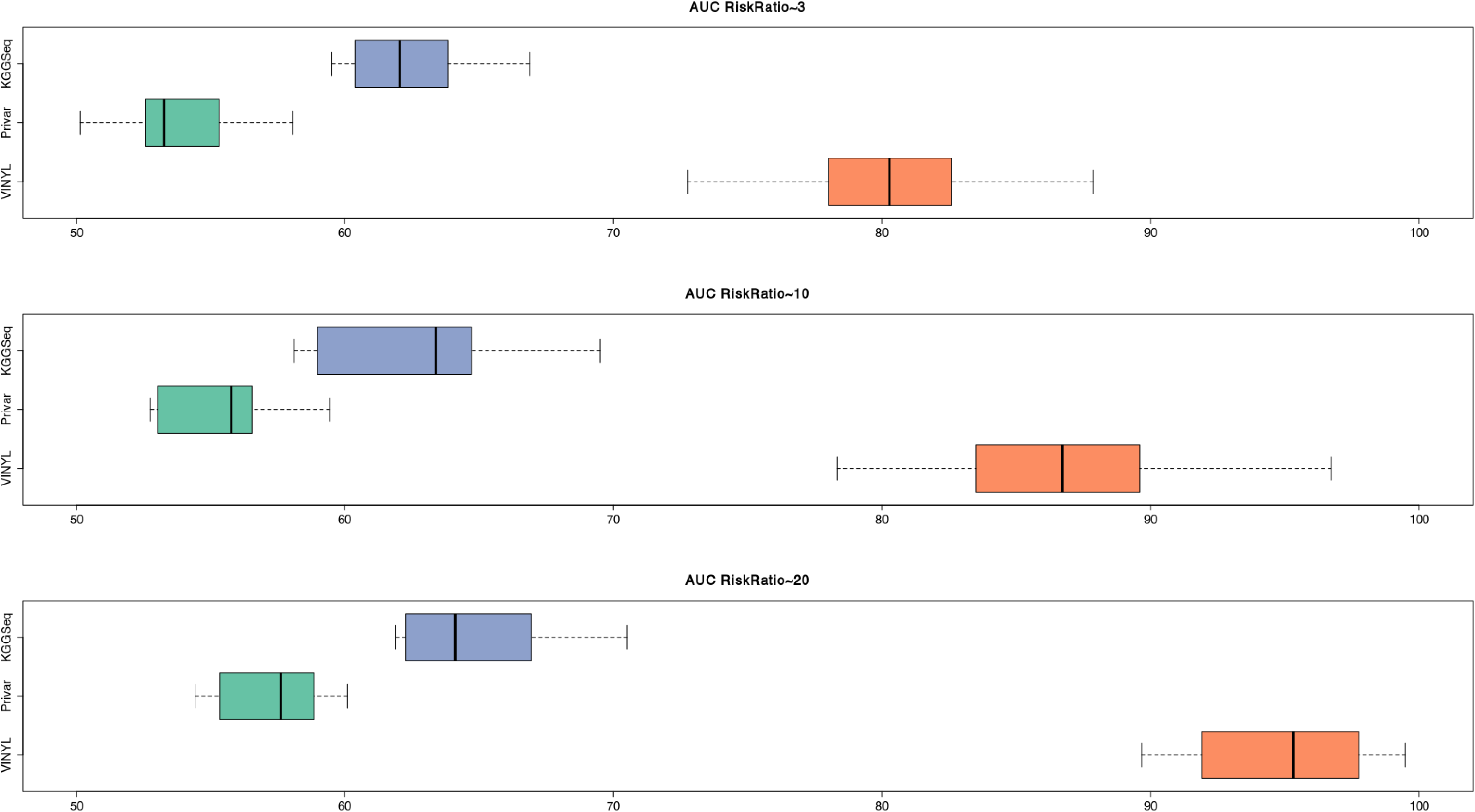
Boxplot of AUC values. Distribution of ROC Area Under the Curve (AUC) values for KGGseq, VINYL and Privar in the detection of simulated pathogenic variants. Distributions of AUC are represented in the form of a boxplot. Panel A, B and C indicate simulations with odd Risk Ratio values of 3, 10 and 20 respectively.

Importantly, we notice that, irrespective of the parameters of the simulations and of the reference control population used, on our simulated dataset VINYL show a consistently higher sensitivity and specificity than any of Privar or KGGseq.

### Evaluation of VINYL on real data

To evaluate the performances of our tool in a realistic scenario, VINYL was applied to a real dataset composed of 38 Italian patients affected by different types of cardiomyopathies, which were previously subjected to genotyping by targeted resequencing of a panel of 115 genes. Expert manual curation of the data resulted in the identification of 27 pathogenic/likely pathogenetic variants (Forleo et al, 2017). Genetic profiles of 107 individuals of Italian ancestry included in the 1000G study (TSI: Toscani in Italia) were used as a control population. Only variants associated with the 115 genes sequenced in Forleo et al. were considered. A total of 53 potentially pathogenic variants (4.02% of the total number) were prioritized by VINYL on this dataset. Notably, all the 27 variants selected by manual curation were recovered, thus achieving perfect accuracy on this dataset (Figure 3A and 3C). On the other hand, only 1 out of 3739 genetic variants in the control population displayed a pathogenicity score higher than the pathogenicity score cut-off value identified by VINYL, suggesting that our method achieves high levels of specificity (Figure 3B). Notably, neither Privar or KGGseq were able to recover the complete collection of the 28 validated and manually curated pathogenic variants (19 and 21 for Privar and KGGseq, respectively) on the same dataset, although the number of variants prioritized by these tools was higher than the number of variants prioritized by VINYL: 84 and 83 respectively for Privar and KGGseq, corresponding to c.a. 6% of the total number of variants in the population of affected individuals. Conversely, both KGGseq and Privar predicted an increased number of pathogenic and/or likely pathogenic variants on the healthy control population: 21 and 22 for Privar and KGGseq respectively, which represent approximately 0.5% of all the variants in the control population (Figure 3B). Taken together. these observations suggest that VINYL achieves higher levels of sensitivity and specificity (Figure 3) also on this dataset.

**Figure 3:**
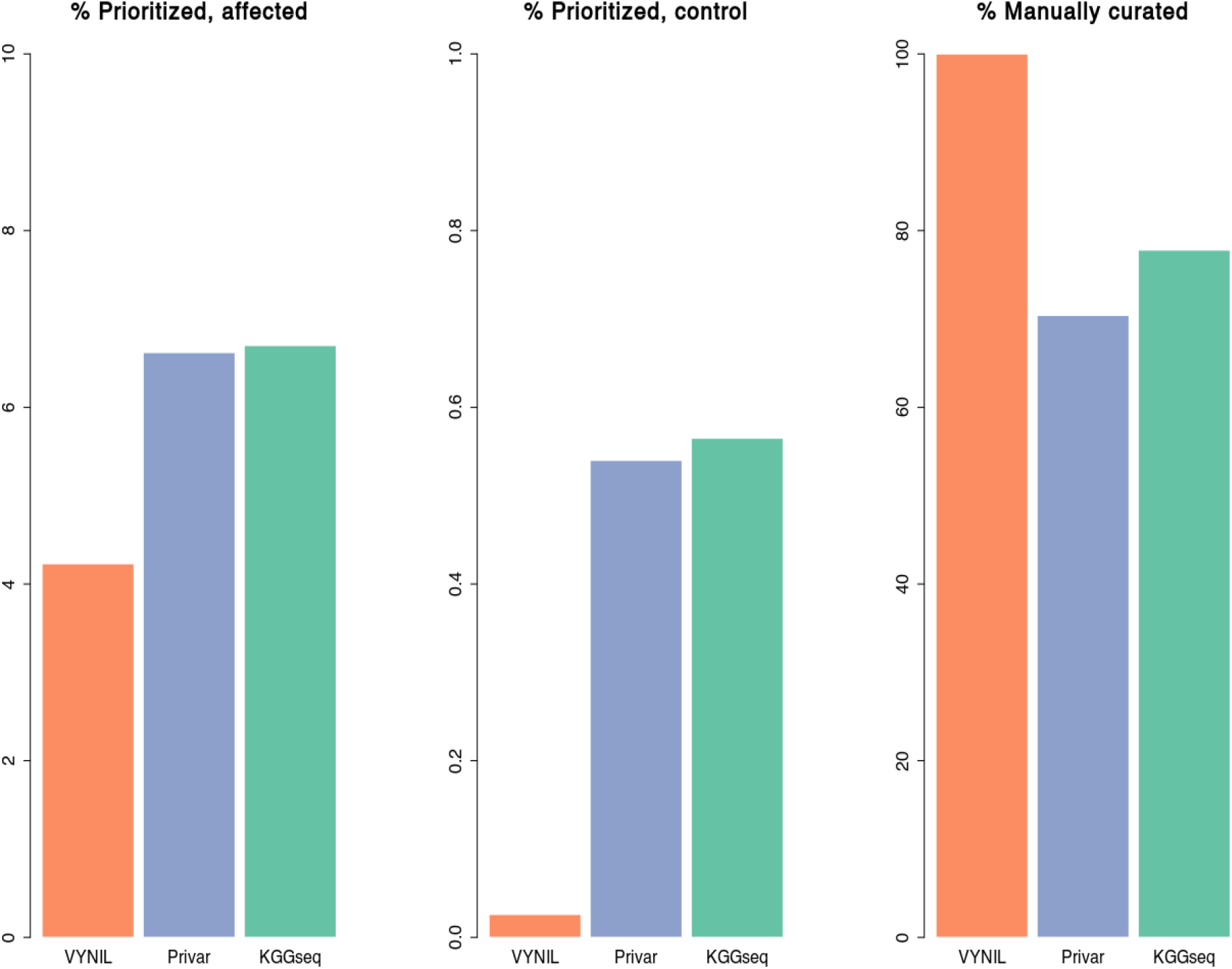
Comparison of VINYL with other start of the art methods for variant prioritization. A) proportion of variants in the population of affected individuals prioritized by each tool. B) Proportion of variants prioritized by each tool in the population of controls. These represent likely false positive calls. C) Proportion of manually curated pathogenic variants according to Forleo et al 2017 recovered by each tool. Orange=VINYL, Blue=Privar, Green=KGGseq.

### Post-processing of the results

Along with tools and resources for variant prioritization, the Galaxy implementation of VINYL incorporates helper applications and utilities to facilitate the post-processing of the data and the interpretation of the results. These include a dimensionality reduction analysis tool, based on Principal Component Analysis (PCA), which can be used for patients stratification, or to identify groups of patients with similar/related disease-associated mutations, and a “burden analysis” utility which can assist in the identification of genes showing a significant increase of pathogenic or likely pathogenic variants. Both utilities produce an explicative graphical output and accept the tabular files generated by VINYL as their main input. An example of the application of these utilities to the Forleo et al dataset is depicted in Figure 4. The PCA analysis displayed in Figure 4A clearly separates controls from affected individuals which split in 2 distinct groups. This could be useful for a better stratification of patients based on profiles of presence/absence of potentially pathogenic variants. As depicted in Figure 4B, the output of VINYL’s burden test analysis consists of a panel where, for every gene, the distribution of VINYL pathogenicity scores, as observed in the cohort of affected individuals, is compared to the corresponding distribution in the control population. A Mann Whitney Wilcoxon test is used to identify genes showing a significant increase in pathogenicity score. Only genes showing a significant p-value are reported in the output. To facilitate a rapid comparison, score distributions are represented in the form of boxplots. Dotted lines are used to indicate the “pathogenicity” cut-off value.

**Figure 4:**
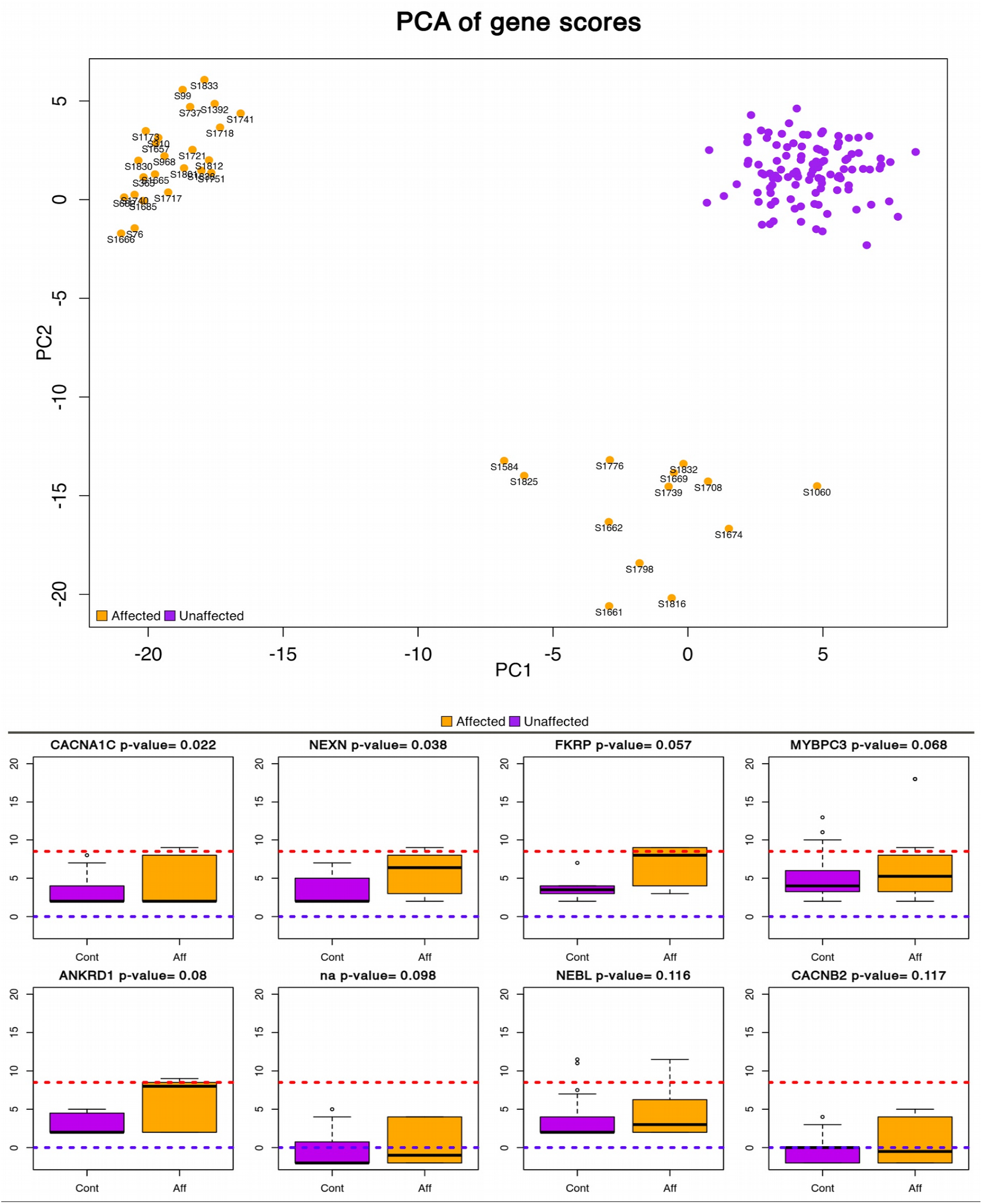
PCA and boxplot analysis of VINYL score. A) Principal component analysis of patients and controls based on VINYL pathogenicity scores. The figure indicates the presence of 2 distinct groups of patients (orange). Both groups of patients are well separated from the control group (purple). B) Comparison of VINYL gene score distribution identifies genes that show a significant increase in pathogenicity score in the population of affected individuals, with respect to the equivalent scores as derived from the control population. Score distributions of each gene are represented in the form of a boxplot. Orange indicates affected individuals, purple indicates controls. Names of the genes and the corresponding Fisher p- value for the increase in score are reported on the top. Dotted red lines indicate the cut-off value used for the identification of pathogenic variants.

## Discussion

The application of genome sequencing technologies to clinical practice is promising a major breakthrough in clinical sciences. However, the systematic integration of genomics into clinical applications poses several challenges, most of which remain unresolved at present. Among these, the systematic functional annotation and interpretation of genomic variants for the rapid identification of genetic traits that might be linked to a pathological condition, a process which is commonly referred to as “variant prioritization” is certainly one of the most critical.

Indeed, this process constitutes one of the major bottlenecks for the rapid analysis of the data, as manual curation by an expert clinician is normally required. Moreover, since different operators/researchers tend to use slightly different methods, criteria, and resources for the functional annotation of genetic variants, the results of these analyses are not always comparable or reproducible, a consideration that limits the possibility to integrate and compare large dataset in a systematic manner and can constitute an issue when the health of patients is at stake.

Here we introduce VINYL, a fully automated system that provides access to a wealth of resources and databases for the functional annotation, and more important to a highly accurate and reproducible method for the prioritization of pathogenic and potentially pathogenic genetic variants. The variant prioritization strategy adopted by VINYL is directly derived from guidelines and best practices that are normally applied in large scale clinical studies and consists in the computation of a composite score that combines different sources of annotation. VINYL extends this approach by using a rapid and rigorous method for the optimization of the scoring system and the selection of a threshold for the identification of potentially pathogenic variants based on the simple observation that a cohort of affected individuals should show an increased number of potentially pathogenic genetic variants when compared to a population of healthy individuals. The method is designed to guarantee high levels of flexibility and permits the incorporation of different types of annotations and resources by the user. By building on the popular Galaxy workflow manager, VINYL is accessible through a simple yet powerful web interface, which enables collaborative work and facilitates the reproducibility of bioinformatics analyses. Encrypted data volumes are used to ensure high levels of data protection. Extensive simulations and analysis of a real dataset, suggest that the approach adopted by VINYL achieves high levels of sensitivity and specificity in different experimental conditions, and more important that our method outperforms currently available state of the art tools in all the conditions herein tested. Although the requirement of a genetically homogeneous “control” population, and the need for a relatively large cohort of affected individuals, limit the applicability of VINYL in cases where only a very limited number of samples is available (see for example analyses of TRIOs or of single patients), we believe that the approach adopted by VINYL is well grounded. Furthermore, this approach is bound to increase its performances over time as it will greatly benefit from the growing number of publicly available data that are being deposited in dedicated databases of genotype-phenotype association such as dbGAP (Mailman et al 2007) and EGA (Lappalainen et al 2015). Indeed, the availability of more data will help in the construction of more accurate scoring systems for specific diseases, which in turn could become applicable also to the analysis of single samples.

Taken together, we believe that, in the light of the results presented in the current study, VINYL will represent a valuable resource to assist in the annotation and prioritization of genetic variants in clinical studies.

## Availability

VINYL is available at http://beaconlab.it/VINYL.

## Acknowledgments

We thank ELIXIR-IIB and ReCaS-Bari for providing computational facilities and Annarita Armenise for technical assistance.

## Funding

This work was supported by the Italian Ministero dell’Istruzione, Università e Ricerca (MIUR): PRIN 2017 and CNRbiomics (PIR01_00017); H2020 Projects ELIXIR-EXCELERATE, EOSC-

Life, and EOSC-Pillar, and Elixir-IIB.

