## Supplemental material for "VINYL: Variant prIoritizatioN bY survivaL analysis"

### Supplementary Table Legends

**Supplementary Table 1:** List of resources for the annotation of genetic variants included in VINYL. First column_ name of the resource according to Annovar. Second column: brief description of the resource. Third column: type of Annovar operation associated with the resource.

| **Resource id** | **Description** | **Annovar operation** |
| --- | --- | --- |
| refGene | Annotation of human genes according to refseq release 80. | gene_ann |
| dbnsfp35a | Prediction of the effect of non-synonymous variants according to several tools, as incorporated in the dbnsfp database | filter |
| esp6500siv2_ea | Allele frequency of genetic variants according to the NHLBI GO Exome Sequencing Project (ESP) | filter |
| exac03nontcga | Allele frequency of genetic variants according to the exac study. Data from the Cancer Genome Atlas (TCGA) samples not included | filter |
| gnomad211_exome, gnomad30_genome | Allele frequency of genetic variants according to the gnomad study. | filter |
| kaviar_20150923 | Allele frequency of genetic variants according to the Kaviar Genomic Variant Database | filter |
| avsnp150 | Allele frequency of genetic variants according to the dbSNP database | filter |
| clinvar_20190305 | Annotation of genetic variants associated with human pathological conditions in the Clinvar database | filter |
| hrcr1 | Allele frequency of genetic variants according to the Haplotype Reference Consortium database. | filter |
| abraom | Allele frequency of genetic variants according to the Brazilian genomic variants (ABRAOM) study | filter |
| AFR.sites.2015_08 | Allele frequency of genetic variants in African populations according to the 1000G study | filter |
| ALL.sites.2015_08 | Allele frequency of genetic variants according to the 1000G study | filter |
| AMR.sites.2015_08 | Allele frequency of genetic variants in American populations according to the 1000G study | filter |
| EAS.sites.2015_08 | Allele frequency of genetic variants in East Asian populations according to the 1000G study | filter |

| | | |
|---|---|---|
| EUR.sites.2015_08 | Allele frequency of genetic variants in European populations according to the 1000G study | filter |
| SAS.sites.2015_08 | Allele frequency of genetic variants in South Asian populations according to the 1000G study | filter |
| dbscsnv11 | Prediction of the effect of genetic variants on splice sites according to dbscSNV version 1.1 both by AdaBoost and Random Forest | filter |
| ENSTFBS | Transcription factor binding sites according to the ENSEMBL regulatory build. For every TF only the sites in the top 90% of the score distribution are reported | region |
| ENSEMBLReg | Segmentation of regulatory genomic elements according to the Ensembl regulatory build | region |
| ENSmiRNA | miRNA target sites according to the Ensembl regulatory build | region |
| gme | Greater Middle East Variome,database of allele frequency of Middle East human populations | |
| GWAS | genetic variants implicated in regions associated to phenotypic traits by one or more Genome-Wide Association Study according to the ENSEMBL database | filter |
| nCER95, nCER90 | genomic regions (10 bp intervals) with an nCER (non-coding Essential Regulation) score higher than 95 and 90 respectively. | region |
| ORegAnno_REGULATORY_P | Polimorphisms with known regulatory effects according to ORegAnno | filter |
| ORegAnno_REGULATORY_R | Segmentation of regulatory genomic elements according to ORegAnno | region |
| ORegAnno_REGULATORY_TFBS | Annotation of Transcription Factor Binding sites according to ORegAnno | region |
| ORegmiRNA | miRNA target sites according to the ORegAnno | region |
| **Annotation of variants associated with eQTL according to the GTEx study.** | | |
| Adipose_Subcutaneous | eQTL | filter |
| Adipose_Visceral_Omentum | eQTL | filter |
| Adrenal_Gland | eQTL | filter |
| Artery_Aorta | eQTL | filter |
| Artery_Coronary | eQTL | filter |
| Artery_Tibial | eQTL | filter |

| | | |
|---|---|---|
| Brain_Amygdala | eQTL | filter |
| Brain_Anterior_cingulate_cortex_BA24 | eQTL | filter |
| Brain_Caudate_basal_ganglia | eQTL | filter |
| Brain_Cerebellar_Hemisphere | eQTL | filter |
| Brain_Cerebellum | eQTL | filter |
| Brain_Cortex | eQTL | filter |
| Brain_Frontal_Cortex_BA9 | eQTL | filter |
| Brain_Hippocampus | eQTL | filter |
| Brain_Hypothalamus | eQTL | filter |
| Brain_Nucleus_accumbens_basal_ganglia | eQTL | filter |
| Brain_Spinal_cord_cervical_c-1 | eQTL | filter |
| Brain_Substantia_nigra | eQTL | filter |
| Breast_Mammary_Tissue | eQTL | filter |
| Cells_EBV-transformed_lymphocytes | eQTL | filter |
| Cells_Transformed_fibroblasts | eQTL | filter |
| Colon_Sigmoid | eQTL | filter |
| Colon_Transverse | eQTL | filter |
| Esophagus_Gastroesophageal_Junction | eQTL | filter |
| Esophagus_Mucosa | eQTL | filter |
| Esophagus_Muscularis | eQTL | filter |
| Heart_Atrial_Appendage | eQTL | filter |
| Heart_Left_Ventricle | eQTL | filter |
| Liver | eQTL | filter |
| Lung | eQTL | filter |
| Minor_Salivary_Gland | eQTL | filter |
| Muscle_Skeletal | eQTL | filter |
| Nerve_Tibial | eQTL | filter |
| Ovary | eQTL | filter |

| | | |
|---|---|---|
| Pancreas | eQTL | filter |
| Pituitary | eQTL | filter |
| Prostate | eQTL | filter |
| Skin_Not_Sun_Exposed_Suprapubic | eQTL | filter |
| Skin_Sun_Exposed_Lower_leg | eQTL | filter |
| Small_Intestine_Terminal_Ileum | eQTL | filter |
| Spleen | eQTL | filter |
| Stomach | eQTL | filter |
| Testis | eQTL | filter |
| Thyroid | eQTL | filter |
| Uterus | eQTL | filter |
| Vagina | eQTL | filter |
| Whole_Blood | eQTL | filter |

**Supplementary Table 2: Classes of functional annotation resources currently recognized by VINYL.** First column: symbol used to represent different types of functional annotation resources. These symbols are used in the VINYL keyword configuration file (kfile) to specify the resource of that type that should be considered for the computation of the score. Second column: brief description of the class of resources. Third column: list of resources of each type currently available in VINYL.

| **Symbol** | **Description** | **Available Resources** |
|---|---|---|
| AF | Publicly available resources of human genotypic variation. Used to infer the Minor Allele Frequency of the variants. When multiple resources are specified, the maximum value is considered | gme,abraom,hrcr1, exac03nontcga,gnomad211_exome, gnomad30_genome kaviar_20150923,avsnp150,esp6500siv2_ea,1000G |
| Effect | Annovar annotation used to derive the predicted functional effect of variants. Normally set to “Func.refGene” and “ExonicFunc.refGene” | NA |
| Dis_DB | Databases of known pathogenic variants to be considered in the computation of the score | Clinvar_20190305 |
| tfbs | TFBS annotations to be considered in the computation of the score | ENSTFBS, ORegAnno_REGULATORY_TFBS |
| Reg | Annotations of regulatory regions to be considered in the computation of the score | ENSEMBLReg, ORegAnno_REGULATORY_R, |

| | | |
|---|---|---|
| | | nCER95, nCER90 |
| NStool | Tools for the evaluation of the effect of non-synonymous variants to be considered in the computation of the score | dbnsfp35a, which currently includes: SIFT, Polyphen2, CADD ,DANN ,MetaLR ,MetaSVM, Provean, VEST, fathmm-MKL and VEST3 |
| mirna | Annotations of miRNA target sites to be included in the score computation | ENSmiRNA, ORegmiRNA |
| gwas | Resources for the annotation of genetic variants associated with phenotypic traits according to A GWAS study | GWAS |
| Splice | Tools for the prediction of the impact of genetic variants on splice sites | dbscSNV_ADA_SCORE, dbscSNV_RF_SCORE |
| RegFilter | List of polymorphisms with known regulatory effects | ORegAnno_REGULATORY_P |
| eQTL | List of tissues (names according to the GTEx) study to be considered for the evaluation of eQTLs | >40, See table S1 |

**Supplementary Table3: Sensitivity and specificity of VINYL on simulated data, using distantly relared control populations**. Levels of sensitivity and specificity of VINYL on simulated data by using different distantly related human geographic population as a control. A) Dataset with 1000 polymorphic sites. B) Dataset with 5000 polymorphic sites. C) Dataset with 10000 polymorphic sites. For A,B, and C size of the simulated cohorts (25,50 or 100 individuals) is reported in the first column. Geographic human populations are indicated in the second column. Columns 3 to 4, 5 to 6 and 7 to 8, report the levels of sensitivity and specificity for identification of simulated pathogenic variants with an odd Risk Ratio of 3, 10 and 20 respectively.

| **1000** | | | | | | | |
|---|---|---|---|---|---|---|---|
| **25 vs 25** | | *RiskRatio~3* | | *RiskRatio~10* | | *RiskRatio~20* | |
| | | *Sens* | *Spec* | *Sens* | *Spec* | *Sens* | *Spec* |
| **25** | *CEU* | 71,66 | 99,43 | 75,93 | 99,38 | 87,06 | 99,62 |
| | *CHB* | 70,93 | 99,22 | 73,52 | 99,51 | 87,23 | 99,46 |
| | *ESN* | 66,38 | 99,19 | 72,71 | 99,22 | 84,26 | 99,52 |
| **50** | *CEU* | 76,47 | 99,36 | 84,67 | 99,42 | 93,19 | 99,14 |
| | *CHB* | 73,72 | 99,53 | 81,85 | 99,58 | 91,46 | 99,30 |
| | *ESN* | 71,04 | 99,38 | 78,28 | 99,52 | 90,76 | 99,17 |
| **100** | *CEU* | 81,83 | 99,16 | 88,58 | 99,60 | 92,02 | 99,30 |
| | *CHB* | 78,29 | 99,15 | 87,59 | 99,59 | 90,75 | 99,16 |
| | *ESN* | 75,14 | 99,57 | 84,05 | 99,39 | 89,28 | 99,36 |
| **5000** | | | | | | | |
| **50 vs 50** | | *RiskRatio~3* | | *RiskRatio~10* | | *RiskRatio~20* | |
| | | *Sens* | *Spec* | *Sens* | *Spec* | *Sens* | *Spec* |
| **25** | *CEU* | 73,97 | 99,34 | 80,01 | 99,34 | 89,11 | 99,42 |
| | *CHB* | 71,62 | 99,40 | 80,65 | 99,54 | 86,30 | 99,57 |
| | *ESN* | 70,71 | 99,55 | 75,34 | 99,14 | 86,16 | 99,26 |
| **50** | *CEU* | 78,66 | 99,43 | 84,75 | 99,57 | 93,84 | 99,44 |
| | *CHB* | 75,49 | 99,26 | 85,00 | 99,17 | 91,16 | 99,61 |
| | *ESN* | 75,78 | 99,54 | 79,95 | 99,36 | 90,13 | 99,36 |
| **100** | *CEU* | 81,60 | 99,53 | 91,35 | 99,50 | 96,85 | 99,56 |
| | *CHB* | 79,69 | 99,59 | 88,96 | 99,15 | 94,95 | 99,43 |
| | *ESN* | 81,13 | 99,45 | 87,43 | 99,50 | 93,06 | 99,49 |
| **10000** | | | | | | | |
| **100 vs 100** | | *RiskRatio~3* | | *RiskRatio~10* | | *RiskRatio~20* | |
| | | *Sens* | *Spec* | *Sens* | *Spec* | *Sens* | *Spec* |
| **25** | *CEU* | 75,98 | 99,12 | 81,97 | 99,35 | 89,27 | 99,53 |
| | *CHB* | 73,51 | 99,17 | 80,37 | 99,44 | 87,37 | 99,47 |
| | *ESN* | 71,37 | 99,29 | 79,51 | 99,16 | 85,48 | 99,19 |
| **50** | *CEU* | 81,09 | 99,20 | 87,21 | 99,60 | 96,54 | 99,34 |
| | *CHB* | 78,98 | 99,48 | 84,31 | 99,41 | 93,74 | 99,21 |
| | *ESN* | 77,01 | 99,28 | 82,24 | 99,31 | 92,65 | 99,38 |
| **100** | *CEU* | 86,96 | 99,31 | 95,19 | 99,40 | 97,33 | 99,19 |
| | *CHB* | 85,47 | 99,42 | 93,30 | 99,16 | 95,48 | 99,29 |
| | *ESN* | 83,16 | 99,29 | 92,22 | 99,44 | 94,38 | 99,41 |

**Supplementary Table4: Sensitivity and specificity of VINYL on simulated data, using heterogenous control populations, with the exclusion of population biased alleles**. Levels of sensitivity and specificity of VINYL on simulated data by using different distantly related human geographic population as a control, with the exclusion of population biased alleles. A) Dataset with 1000 polymorphic sites. B) Dataset with 5000 polymorphic sites. C) Dataset with 10000 polymorphic sites. For A,B, and C size of the simulated cohorts (25,50 or 100 individuals) is reported in the first column. Geographic human populations are indicated in the second column. Columns 3 to 4, 5 to 6 and 7 to 8, report the levels of sensitivity and specificity for identification of simulated pathogenic variants with an odd Risk Ratio of 3, 10 and 20 respectively.

| **1000** | | | | | | | |
|---|---|---|---|---|---|---|---|
| **25 vs 25** | | *RiskRatio~3* | | *RiskRatio~10* | | *RiskRatio~20* | |
| | | *Sens* | *Spec* | *Sens* | *Spec* | *Sens* | *Spec* |
| **25** | *CEU* | 72,01 | 99,39 | 77,52 | 99,54 | 88,61 | 99,58 |
| | *CHB* | 70,69 | 99,57 | 77,41 | 99,61 | 88,09 | 99,53 |
| | *ESN* | 71,32 | 99,43 | 76,49 | 99,46 | 87,67 | 99,46 |
| **50** | *CEU* | 78,09 | 99,32 | 85,52 | 99,38 | 93,25 | 99,49 |
| | *CHB* | 76,65 | 99,29 | 84,51 | 99,42 | 93,09 | 99,52 |
| | *ESN* | 75,84 | 99,56 | 84,59 | 99,31 | 90,60 | 99,62 |
| **100** | *CEU* | 81,48 | 99,25 | 89,33 | 99,42 | 94,91 | 99,40 |
| | *CHB* | 81,35 | 99,32 | 88,54 | 99,62 | 95,25 | 99,52 |
| | *ESN* | 80,91 | 99,41 | 88,18 | 99,41 | 93,72 | 99,63 |
| **5000** | | | | | | | |
| **50 vs 50** | | *RiskRatio~3* | | *RiskRatio~10* | | *RiskRatio~20* | |
| | | *Sens* | *Spec* | *Sens* | *Spec* | *Sens* | *Spec* |
| **25** | *CEU* | 76,36 | 99,68 | 81,76 | 99,27 | 90,09 | 99,50 |
| | *CHB* | 75,45 | 99,58 | 80,99 | 99,48 | 90,10 | 99,65 |
| | *ESN* | 74,74 | 99,48 | 79,14 | 99,48 | 89,43 | 99,49 |
| **50** | *CEU* | 79,48 | 99,52 | 86,28 | 99,64 | 95,40 | 99,68 |
| | *CHB* | 78,79 | 99,41 | 85,18 | 99,49 | 94,28 | 99,24 |
| | *ESN* | 78,68 | 99,34 | 84,70 | 99,67 | 92,40 | 99,33 |
| **100** | *CEU* | 83,27 | 99,65 | 92,71 | 99,23 | 98,49 | 99,46 |
| | *CHB* | 82,83 | 99,56 | 91,29 | 99,46 | 98,64 | 99,46 |
| | *ESN* | 80,66 | 99,46 | 91,22 | 99,18 | 96,39 | 99,68 |
| **10000** | | | | | | | |
| **100 vs 100** | | *RiskRatio~3* | | *RiskRatio~10* | | *RiskRatio~20* | |
| | | *Sens* | *Spec* | *Sens* | *Spec* | *Sens* | *Spec* |
| **25** | *CEU* | 77,02 | 99,19 | 83,22 | 99,60 | 90,85 | 99,61 |
| | *CHB* | 76,48 | 99,26 | 81,53 | 99,17 | 90,17 | 99,50 |
| | *ESN* | 75,10 | 99,68 | 80,63 | 99,65 | 88,35 | 99,61 |
| **50** | *CEU* | 81,62 | 99,55 | 87,27 | 99,33 | 96,92 | 99,17 |
| | *CHB* | 80,54 | 99,67 | 86,53 | 99,66 | 96,99 | 99,58 |
| | *ESN* | 79,71 | 99,27 | 85,37 | 99,42 | 96,68 | 99,30 |
| **100** | *CEU* | 86,94 | 99,18 | 96,49 | 99,26 | 97,96 | 99,39 |
| | *CHB* | 87,07 | 99,45 | 95,88 | 99,31 | 97,50 | 99,26 |
| | *ESN* | 84,42 | 99,68 | 95,02 | 99,31 | 96,92 | 99,50 |

### Configuration files and parameters

VINYL requires several configuration files for the calculation of its pathogenicity score. Although some of these files are not mandatory, it is highly recommended to incorporate into VINYL as much information as possible in order to obtain high levels of sensitivity and accuracy. An example of each of type of configuration file, can be found at in the Galaxy implementation of VINYL, at the following link: http://90.147.75.93/galaxy/library/list#folders/Fb04ca61b52e20534 or in the VINYL Github repository. A description of the configuration files and the input parameters of VINYL is provided in the following section.

VINYL uses different sources of annotation for the computation of its pathogenicity score, users can specify the type and the list of resources to be used by the means of the “keywords” configuration file. This is a configuration file that specifies the keywords that are used by VINYL for the extraction of relevant annotations from VCF files and for the computation of the pathogenicity score. Names of these keywords need to match exactly those used by Annovar (Wang et al 2019). Currently VINYL discriminates between 8 different types of "keywords", which are used to calculate different components of the score. see supplementary Table 1 and supplementary Table 2, for a complete description of the resources available in VINYL.

- **AF keywords**: specify the databases/resources that are used to obtain Allele Frequency annotations. When multiple resources are provided the maximum value is considered
- **NStool keywords:** specify the tools for the evaluation of the functional effects of Non Synonymous variants that are considered by VINYL for the computation of the Disruptive non synonymous variants score. More than one tool can be specified.
- **Effect keywords:** these keywords set the columns that are used by VINYL to extract the predicted annotation of the functional effects of the variants. If values associated with these keys match any of the deleterious effects as listed in the functional effects file (see below) the pathogenicity score is incremented
- **Splice keywords:** specify the databases/resources that are used to evaluate the effects of genetic variants on splice site. In the current implementation of VINYL these correspond with the dbscSNV_ADA_SCORE and the dbscSNV_RF_SCORE. Both scores are derived from the dbscSNV database (Jian et al, 2014)

- **Reg keywords**: specify the databases to be considered for the annotation of regulatory regions. Defaults to ORegAnno_REGULATORY_R and ENSEMBLReg. The nCER90 and nCER95 databases can be specified as well.
- **tfbs keywords:** specify the databases to be considered for the annotation of Transcription Factor Binding sites regions. Defaults to ORegAnno_REGULATORY_TFBS and ENSTFBS
- **mirna keywords:** specify the databases to be considered for the annotation of miRNA binding sites. Defaults to ENSmiRNA and ORegmiRNA
- **gwas keywords:** these keywords are used to indicate the resources to be used for the annotation of SNPs implicated with phenotypic traits of interest as specified in the "symptoms" configuration file (see below) according to a Genome Wide Association Study. At the time being, the annovar database incorporated in VINYL includes only one resource for this type of annotation, the GWAS reference database, as obtained from https://www.ebi.ac.uk/gwas/https://www.ebi.ac.uk/gwas/ (Buniello et al, 2019)

The format of the keyword file is very simple: this a plain tabular file where each keyword (be aware that the names should match exactly the names used by Annovar) is followed by a qualification that specifies its type. As outlined above types can be any of: AF, Effect, NStool, Spliece, Reg, tfbs, mirna, and gwas.

Predicted functional effects which should be considered deleterious, can be specified by the means of the functional effects configuration file (efile). The file has a very simple format: each functional effect is provided in a separate line. The convention used to indicates these effects is the same as that applied by Annovar (Wang et al 2010). Please refer to the Annovar documentation for a thorough explanation of the annotation of predicted functional effects by Annovar.

Symptoms of the pathological condition under study, can be specified by the symptoms configuration file. This file provides a list of symptoms or related keywords that are used by VINYL to screen Clinvar (Laundrum et al, 2014) Annotations and identify variants that have been implicated in similar pathologies or phenotypes. The file is provided in a simple text format, where each of the keyword needs to be reported on a separate line. See the following link: http://90.147.75.93/galaxy/library/list#folders/Fb04ca61b52e20534/datasets/d7c24196f42c4cea for an example of a valid file. Users are encouraged to check the controlled vocabulary of names and symptoms associated with the pathological condition of interest in the OMIM database (Amberger et al, 2019).

A list of genes previously implicated in the pathological condition under study can be specified via the gene list configuration file (gfile). The file is in simple text format, with each gene reported on a separate line. Genes are indicated by their official gene symbol. An example is provided at the following link: http://90.147.75.93/galaxy/library/list#folders/Fb04ca61b52e20534/datasets/ac319686878db63c

List of tissues to be considered for the analysis/annotation of expression Quantitative Trait Loci (eQTL) can be specified through the "eQTL" configuration file. Names of tissues need match names used in the GTEx project (GTEx Consortium, 2013). Again, this is simple text file with the name of the tissues in a separate line.

See http://90.147.75.93/galaxy/library/list#folders/Fb04ca61b52e20534/datasets/37daec133973324a for an example.

Users are also required to provide additional parameters in the form of simple text entries or numbers for cut-off values used in the computation of the pathogenicity score, these include:

- **Disease name:** Name or functional description of the pathological condition. This parameter is used to perform a soft check of the annotation in Clinvar and in GWAS studies to identify variants that have been previously implicated in the disease. Clinvar or GWAS annotations that match the value provided by this parameter and/or of the disease, as specified in the sfile, are considered for the computation of the pathogenicity score. This parameter is **highly recommended**. Users are encouraged to check the controlled vocabulary of names and symptoms associated with the pathological condition of interest in the OMIM database (Amberger et al, 2019). If the patients are affected by different pathological conditions, multiple names can be provided, separated by a "#". For example: "cardiomyopathy#dilated" specifies the keywords "cardiomyopathy" and "dilated".
- **Over-representation score cut off:** Cut off value used for computation of the over-representation score. The increment value specified by OR (see main text) is added to the pathogenicity score only for variants that have an allele count equal to or greater than this cut off. As a rule of thumb, this should be set to approximately 5-10% of the size of your cohort of individuals. Defaults to 10% of the size of the cohort
- **Allele frequency cutoff:** Cut off value for the allele frequency cut-off used in the computation of the RV score. The value specified by the RV score is added to the pathogenicity score only for

variants that show an allele frequency lower or equal to this cut-off value

- **Hereditary model of the disease:** Currently VINYL supports 3 (mutually exclusive) types of disease model: Autosomic Dominant, Autosomic Recessive and X-linked. Disease models can be specified by the means of the AD (Autosomic Dominant) and by the XL (X-Linked) parameters. These parameters accept logical values: T=TRUE, F=FALSE.
